# Ok-*k*-mer: Selecting *k*-mer length ensembles for calculating genomic distance

**DOI:** 10.64898/2026.11.06.764926

**Authors:** Stephen Hwang, Ben Langmead

## Abstract

Current methods for computing sequence similarity across large groups of genome sequences tend to do so at the level of ***k***-mers since this is far more efficient than base-by-base alignment. While these methods use default ***k***-mer lengths, recent work shows that the particular choice of ***k***-mer length greatly affects accuracy, and that using multiple values of ***k*** can boost accuracy relative to the best single choice of ***k***, particularly when the goal is to estimate genomic distances that reflect true phylogenetic relationships. Here, we introduce Ok-***k***-mer, a tool to systematically select weighted ***k***-mer length ensembles for pairwise genomic distance estimation, thereby increasing the accuracy of those estimates. Given a phylogenetic tree, Ok-***k***-mer ranks genome pairs by Jaccard distance across a range of ***k*** and uses a progressive search to maximize the Spearman correlation, ***ρ***, between weighted-average ***k***-mer ranks and branch distances. Across eight Genome Taxonomy Database datasets, Ok-***k***-mer ensembles improved correlation over standard default ***k***-mer lengths. In the Chlamydiales taxonomy, a three-length ensemble ***k* = (9, 17, 22)** achieved a 23.9% relative improvement over the best single length (***k* = 20**) and a 36.7% improvement over a commonly used default of **31**-mers. We demonstrate that Ok-***k***-mer ***k***-mer length ensembles remain stable (i.e. lengths and weights change only moderately) under genome subsampling within a taxonomy but vary across datasets. Ok-***k***-mer can annotate all subtrees of a phylogeny, which reveals distinct patterns of ensembles reflecting potential differences in genome evolution. Ok-***k***-mer provides a framework for moving beyond a “one-***k***-fits-all” ***k***-mer analysis, improving genomic distance estimation in the era of massive-scale genomics.

## 1 Introduction

The rapid growth of our reference genome collections necessitates accurate and efficient methods for comparing whole genome sequences. As a computational expedient, many such methods compare genomes using *k*-mers, substrings of a fixed length *k*, and avoid performing base-by-base alignments. But *k*-mer-based genomic distance methods have limitations: (1) though they are faster than alignmentbased methods, they are not as accurate [1, 2], and (2) the choice of *k*-mer length impacts accuracy. For example, Nasko et al. [3] showed that as databases grow, *k*-mer-based taxonomic read classifications lose specificity. Nevertheless, widely used tools to compute genomic distances use fixed, default *k*-mer lengths often with little justification (Table A1): Mash uses *k* = 21, sourmash and Skmer use *k* = 31, and Kmer-db uses *k* = 18 [4–7]. FastANI and skani use shorter defaults, *k* = 16 and *k* = 15, respectively, for mapping and comparing homologous regions [8, 9]. While these defaults might function well for specific tasks, a default appropriate for one task or dataset will not be optimal for another.

Past genomics methods have sought to move beyond the use of a single, fixed *k*-mer length. KmerGenie estimates *k*-mer abundance histograms across candidate lengths to select a length for genome assembly [10]. PopPUNK uses comparisons across multiple *k*-mer lengths to estimate divergence in shared sequence and differences in accessory sequence content [11]. Slope-SpaM estimates phylogenetic distances from how the number of word matches changes with word length [12]. MEMO supports arbitrary-length *k*-mer membership and conservation queries from a single pangenome index for *k*-mers of a designated pivot genome [13]. Yet, there are no established tools for selecting or recommending *k*-mer lengths for Jaccard-based genome comparison.

Here, we introduce Ok-*k*-mer, a tool for selecting weighted ensembles of *k*-mer lengths for pairwise genomic distance estimation. Building on the EvANI benchmarking framework [2], Ok-*k*-mer uses a reference phylogeny to select lengths and weights that maximize the Spearman correlation between weighted-average Jaccard distance ranks and reference tree distances. Across eight Genome Taxonomy Database (GTDB) datasets, the selected ensembles improve agreement with phylogenetic distances relative to single *k*-mer lengths and commonly used default lengths. Recommendations vary across datasets but remain stable (i.e. with only moderate changes in the *k*-mers and weights selected) under genome subsampling. By examining Ok-*k*-mer’s results across nodes of a phylogeny, we observe that different decisions are indeed needed for different subtrees. These results provide a basis for datasetspecific length selection in genomic comparisons, with potential applications to genome placement and taxonomic classification.

## 2 Results

### 2.1 Ok-*k*-mer overview

Ok-*k*-mer selects weighted *k*-mer length ensembles for accurate pairwise genomic distance estimation across a phylogenetic tree. Given a phylogenetic tree with branch lengths, Ok-*k*-mer computes the *k*-mer Jaccard distance for all genome pairs across a range of *k* (Fig. 1; Methods 4.2–4.5). Reference distances are used to select a dataset-specific *k*-mer length ensemble from genomes with known relationships. The selected ensemble is then used to calculate pairwise relationships to new genomes related to the reference dataset, such as with genomic placement. Ok-*k*-mer ranks genome pairs by their tree distances (Fig. 1B) and follows the EvANI benchmarking framework [2] for evolutionary distance estimation based on the rank correlation of genome pairs across a phylogenetic tree (Methods 4.2). Ok-*k*-mer establishes a novel progressive search approach to identify the best *k*-mer length ensembles (Fig. 1C). By weighting the evidence from different lengths, Ok-*k*-mer can disambiguate ties between genome pairs and refine the ultimate ranking.

**Fig. 1.**
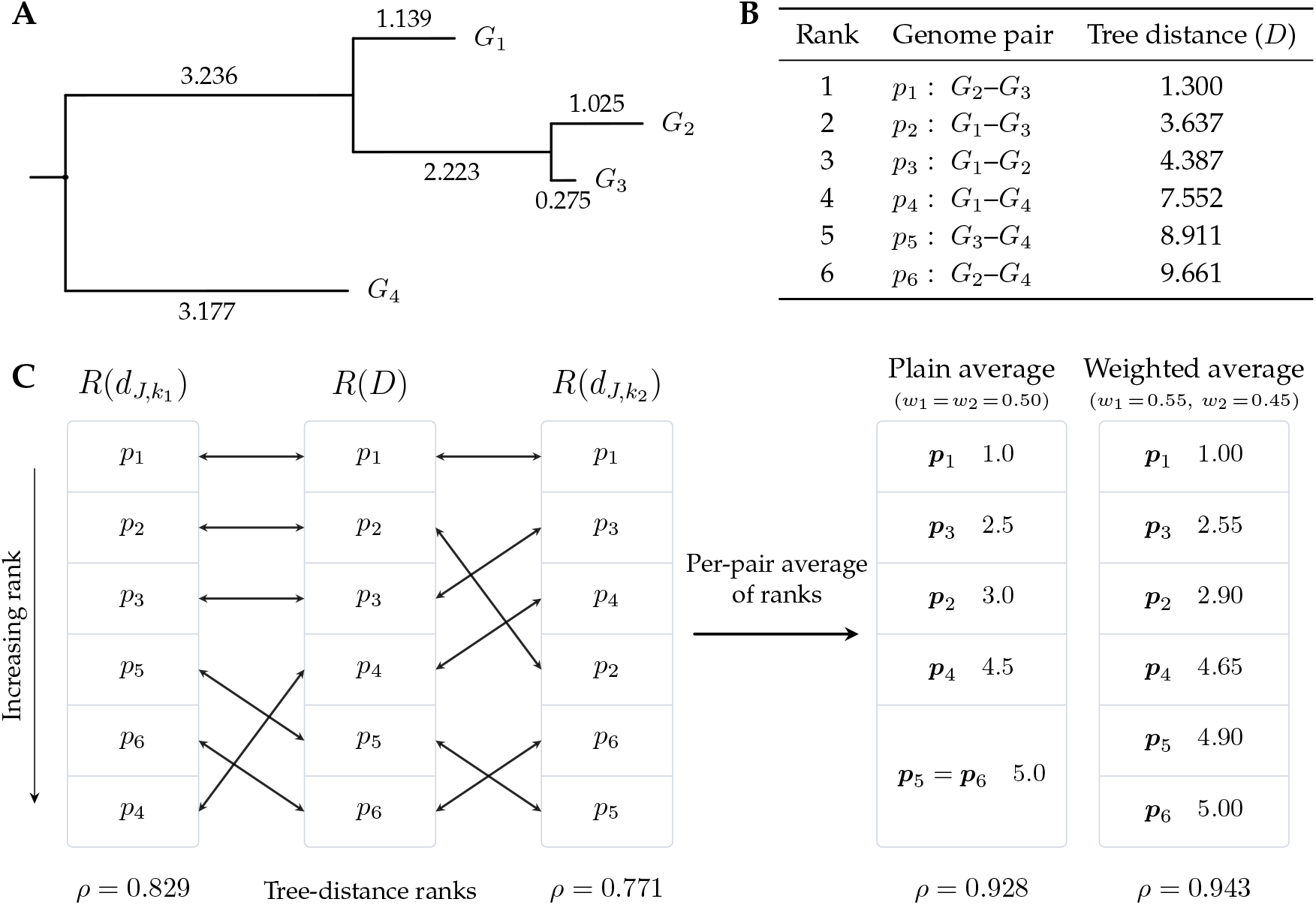
Overview of rank aggregation in Ok-*k*-mer on an example tree. **(A)** Taxonomic tree (*T*) with four genomes (*G*_1_, … , *G*_4_) and annotated branch lengths. **(B)** All pairs of genomes (*p*_*n*_) ranked by their tree distances (*D*) in *T* . **(C)** Rank ordering of Jaccard distances for two *k*-mer lengths (*R*(*d*_*J,k*_)) and the correlation of their per-pair average ranks with the ranked true tree distances (*R*(*D*)). Spearman correlation (*ρ*) is shown below each rank array. The plain average of 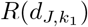 and 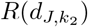 yields a higher Spearman correlation (*ρ* = 0.928) than that obtained from either single-*k* Jaccard distance ranking. The weighted average (*w*_1_ = 0.55, *w*_2_ = 0.45) resolves the *p*_5_ = *p*_6_ tie in the plain average and further improves Spearman correlation (*ρ* = 0.943).

We evaluate eight Genome Taxonomy Database (GTDB) datasets [14] spanning various sizes and taxonomic ranks. We demonstrate that datasets benefit from information from multiple *k*-mer lengths and that no single *k*-mer length is optimal across datasets. Ok-*k*-mer can annotate all internal nodes of a phylogeny and inform genome placement and classification. Ok-*k*-mer captures dataset-specific information; the recommendations are stable across subsets of the same taxonomy, but differ across distinct datasets and across distinct subtrees of the same phylogeny.

### 2.2 *K*-mer length ensembles benefit all-pairs distance estimates

We hypothesized that using well-selected *k*-mer length ensembles could improve the concordance of pairwise distances for all-pairs comparisons in a taxonomic tree. To evaluate this, we used established genomes and accompanying taxonomies with branch lengths. Although complete reference distances are not generally available in practice, they provide a controlled benchmark for determining whether combining *k*-mer length information improves concordance with evolutionary distance. We compared the best *k*-mer length ensembles containing one, two, or three *k*-mer lengths using the global search (Methods 4.5). Across the Chlamydiales taxonomy, *k* = 20 yields the highest correlation among single *k*-mer lengths (*ρ* = 0.4240; Fig. 2A). Combining ranks from two lengths improves concordance, with the optimal pair (*k*_1_, *k*_2_) = (9, 18) achieving *ρ* = 0.5132 (Fig. 2B). We assume that weights can never be zero, since adding zero-weight lengths to an ensemble will not change the result. Using three lengths further improves *ρ*: the optimal ensemble (*k*_1_, *k*_2_, *k*_3_) = (9, 17, 22), with weights (0.45, 0.30, 0.25), achieves *ρ* = 0.5254 (Fig. 2C). Notably, clusters of well-performing *k*-mers are visible in the heatmaps. Although we report the globally optimal *k*-mer length ensembles, a dataset may contain many similarly highperforming *k*-mer combinations that yield *ρ* values that are extremely close to the global maximum. For example, considering just the *k*-mer length ensembles for Chlamydiales, we observed that there were 164 distinct three *k*-mer ensembles that yielded higher *ρ* than the best two *k*-mer length ensemble. We also observed that there were 27 unique three *k*-mer length ensembles that achieved a *ρ* within 0.0025 of the optimal *ρ*.

**Fig. 2.**
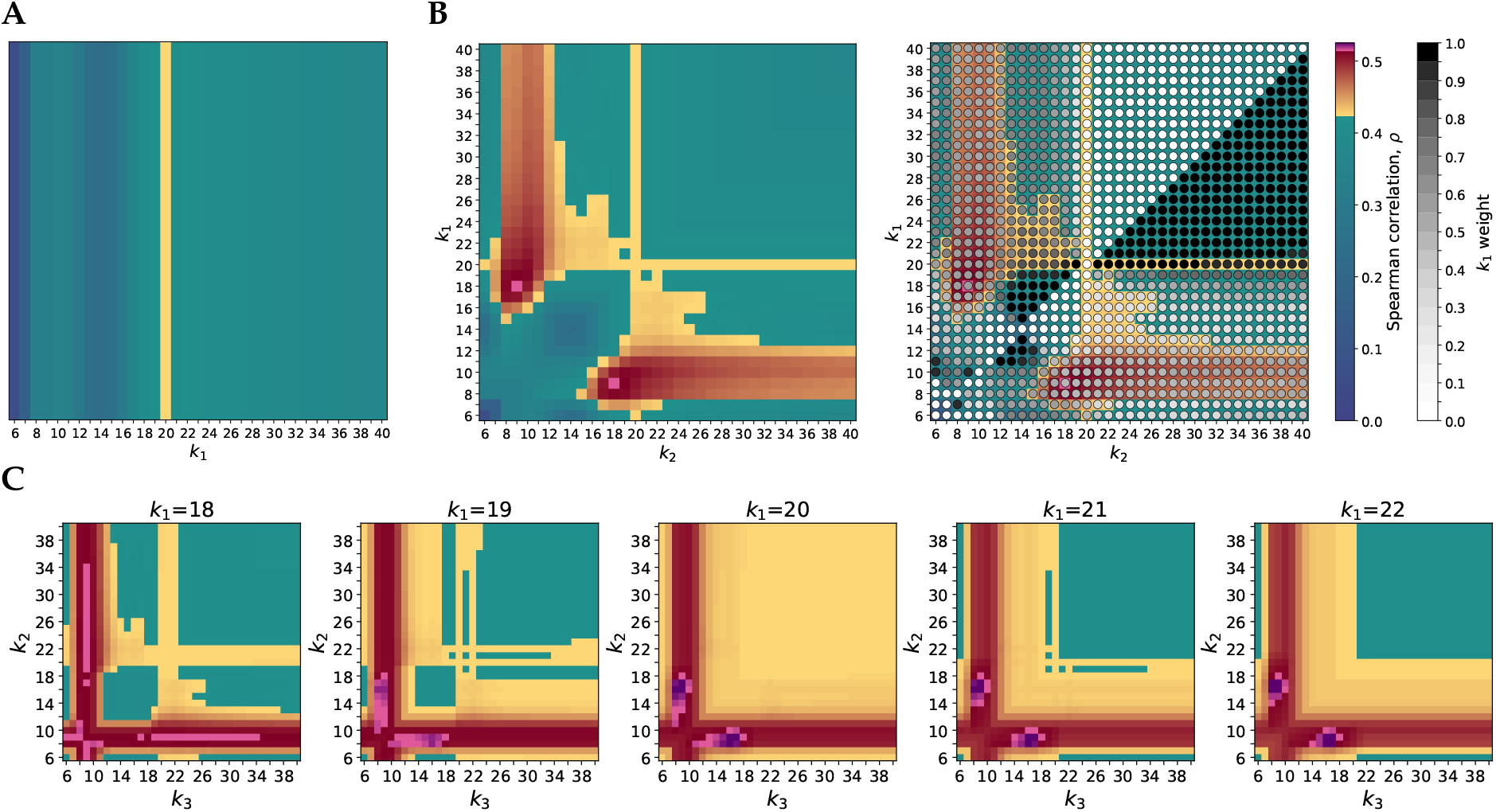
The Chlamydiales subtree of the GTDB taxonomy (*n* = 288) benefits from *k*-mer length ensembles. The legend for Spearman correlation *ρ* is shared across all plots and is shown at right. The legend colors are divided into three ranges (blue-to-green, yellow-to-red, pink-to-purple), with each range adjusted so that the maximum correlation for one *k*-mer tier begins the next color range. **(A)** Color bar showing Spearman correlation values for single *k*-mer lengths with *k* ∈ {6, … , 40} . The yellow stripe marks the highest-performing *k*-mer length of 20 (*ρ* = 0.4240). **(B)** Heatmap showing, for each pair of *k*-mer lengths, the correlation *ρ* obtained from the best weighted average of their ranks (left). Weights *w* ∈ [0, 1] in steps of 0.05 are superimposed on each cell (right). The pink square marks the bestperforming two-*k*-mer ensemble (*ρ* = 0.5132), which is (*k*_1_, *w*_1_) = (9, 0.45) and (*k*_2_, *w*_2_) = (18, 0.55). **(C)** Heatmaps showing the correlation *ρ* for the best weighted average between the ranks of all ensembles of three *k*-mer lengths centered on the best single *k*-mer, *k* = 20. Weights are not depicted in this panel. The darkest purple square marks the best-performing three-*k*-mer ensemble (*ρ* = 0.5254) with (*k*_1_, *w*_1_) = (9, 0.45), (*k*_2_, *w*_2_) = (17, 0.30), and (*k*_3_, *w*_3_) = (22, 0.25).

We applied our global *k*-mer length ensemble search algorithm to eight GTDB datasets (Methods 4.1, 4.5; Table 1). Seven of the datasets benefited from a weighted average of two *k*-mer length ranks. Five of the datasets benefited from a three-*k*-mer-length ensemble. One dataset, Caldisericia, required only a single *k*-mer length. By contrast, the Chlamydiales Spearman correlation improved from 0.4240 to 0.5254 (≈ 24% increase) from using a single *k*-mer to using three well-separated *k*-mers. On the eight GTDB datasets, the default lengths *k* = 21 and *k* = 31 (Table A1) yield Spearman correlations that average 3.70% and 10.33% below the Ok-*k*-mer *ρ*, respectively (Table A4). We search for ensembles of up to three *k*-mer lengths because extending the search to a fourth length increased runtime with only marginal improvements in correlation. For example, in Bacillales A, the best four-length ensemble *k* = (6, 7, 23, 24) increased *ρ* by only 0.00000325 relative to the best three-length ensemble.

**Table 1.**
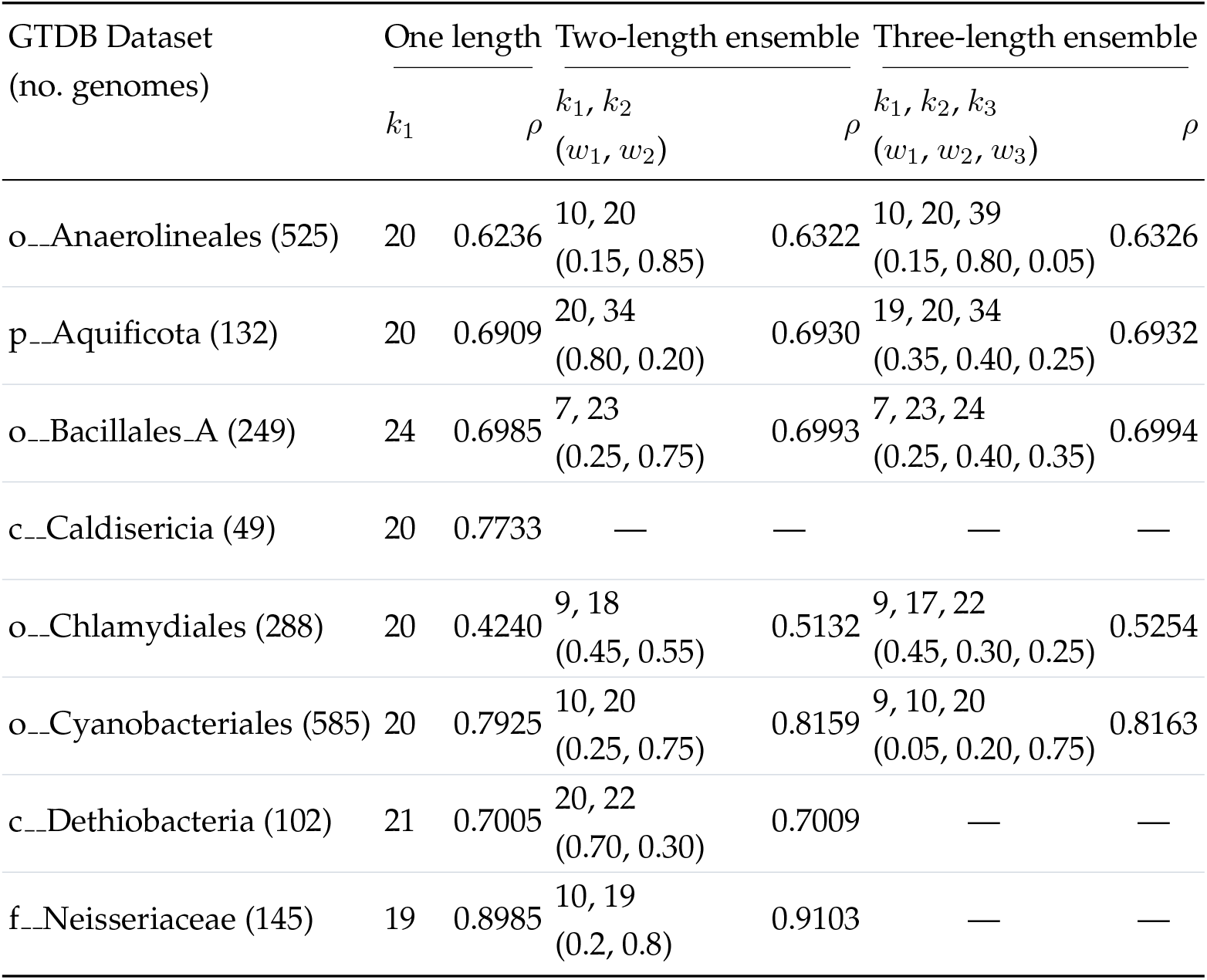
Comparison of the best *k*-mer length ensembles for computing distances across GTDB datasets. Dataset name prefixes p-, c-, o-, and f denote phylum, class, order, and family, respectively. *ρ* is the Spearman correlation coefficient between the weighted *k*-mer rank ensemble and the reference branch distances. For multi-*k* ensembles, weights are shown in parentheses below the selected *k*-mer lengths. The ordering of the weights corresponds to the order of the *k*-mers. Dashes indicate that the correlation is not improved by adding further *k*-mer lengths.

### 2.3 Ok-*k*-mer annotates the subtrees of a phylogenetic tree

Applying Ok-*k*-mer to individual subtrees reveals variation in recommended lengths and weights across the phylogeny (Fig. 3). The internal node annotations reveal different ensembles of *k*-mers across subtrees. For example, some root-to-leaf paths yield nodes with similar *k*-mer recommendations, whereas others yield nodes with varying recommendations (Fig. 3). Adjacent nodes with varying *k*-mer recommendations but similar Spearman *ρ* may represent branches for which several *k*-mer combinations yield close-to-optimal correlations.

**Fig. 3.**
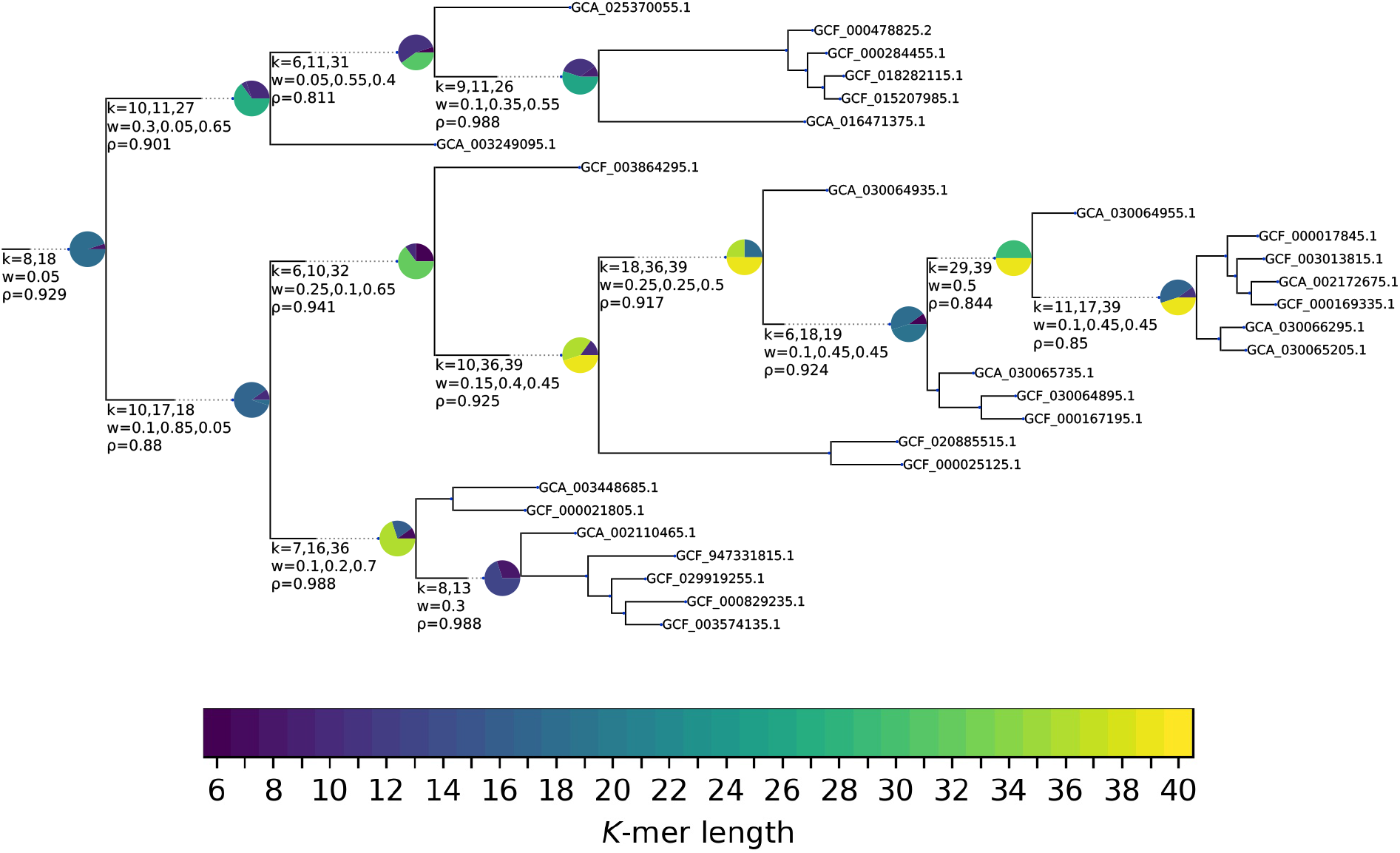
A subtree of the full Cyanobacteriales taxonomy depicting Ok-*k*-mer annotations for optimal *k*- xmer lengths and weights for up to three *k*-mer lengths. Nodes are annotated with text (*k*-mer lengths, weights, and *ρ*) and corresponding pie charts. Slice color represents *k*-mer length, and slice size represents weight. Different branches of the tree require different *k*-mer length ensembles. Some root-to-leaf paths yield nodes with similar ensembles (top branch). Other paths yield nodes with varying *k*-mer recommendations (middle branch). These nodes represent datasets in which a variety of *k*-mer ensembles yield close-to-optimal Spearman *ρ* values.

These recommendations can inform downstream genome placement and taxonomic classification; depending on the complexity of the database and where the query genome falls, Ok-*k*-mer can recommend dataset-dependent *k*-mers for maximal distinguishing power.

### 2.4 Ok-*k*-mer recommendations are stable and dataset-specific

Ok-*k*-mer recommendations are generalizable to larger sets of the same phylogeny. We randomly masked 10%–60% of the genomes in each phylogeny and compared the resulting correlations to that of the original (0% masked) dataset. At each masking percentage, we ran 15 replicates per dataset and plotted the mean and spread (*±* 1 standard deviation) of Spearman *ρ* (Fig. 4). Within each dataset, mean Spearman *ρ* remained close to the full-dataset correlation across masking fractions. Ok-*k*-mer gives similar recommendations to a phylogeny when more genomes are added.

**Fig. 4.**
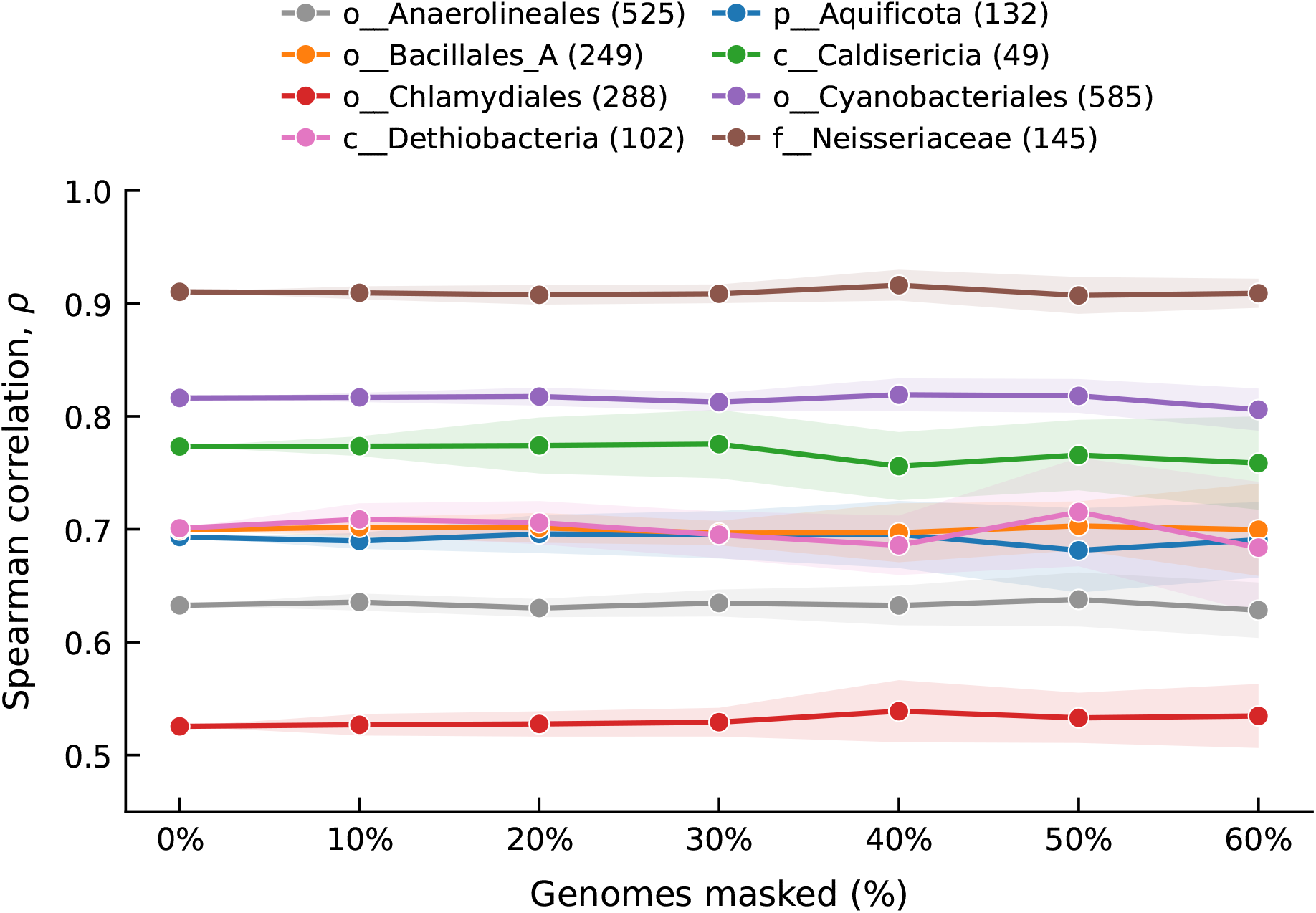
Mean Spearman *ρ* across 15 independent simulations at masking fractions from 0% to 60% demonstrates the stability of Ok-*k*-mer recommendations. The spread is *±*1 standard deviation around the mean *ρ*.

Across 30% masking trials from the same dataset, Ok-*k*-mer yielded similar selected *k*-mers. The first three independent trials at 30% masking corroborate the stability of the resulting correlations under genome subsampling (Table 2). Additionally, the selected lengths differ across datasets, further supporting dataset-specific selection of *k*-mer length ensembles (Fig. A1).

**Table 2.** Ok-*k*-mer recommendations from the first three independent simulation trials after masking 30% of genomes in each GTDB dataset. In parentheses is the number of genomes retained after masking over the full number of genomes. Rows report the selected *k*-mer lengths, their weights, and Spearman *ρ*. Selected *k*-mers and resulting *ρ* are similar across trials for each dataset.

| GTDB Dataset<br>(no. genomes retained) | Trial | Selected *k*-mers | Weights | Spearman ρ |
|---|---|---|---|---|
| o__Anaerolineales (368/525) | 1 | 6, 10, 20 | 0.05, 0.15, 0.80 | 0.6336 |
|  | 2 | 10, 20, 40 | 0.15, 0.80, 0.05 | 0.6132 |
|  | 3 | 10, 20, 22 | 0.15, 0.65, 0.20 | 0.6456 |
| p__Aquificota (93/132) | 1 | 20, 34 | 0.85, 0.15 | 0.7173 |
|  | 2 | 20, 34 | 0.90, 0.10 | 0.6646 |
|  | 3 | 20, 33 | 0.85, 0.15 | 0.6932 |
| o__Bacillales_A (175/249) | 1 | 6, 23, 24 | 0.05, 0.70, 0.25 | 0.7084 |
|  | 2 | 6, 23 | 0.45, 0.55 | 0.6967 |
|  | 3 | 6, 7, 23 | 0.05, 0.25, 0.70 | 0.6960 |
| c__Caldisericia (35/49) | 1 | 19, 20 | 0.10, 0.90 | 0.7261 |
|  | 2 | 20, 21 | 0.90, 0.10 | 0.7742 |
|  | 3 | 20, 21 | 0.40, 0.60 | 0.7518 |
| o__Chlamydiales (202/288) | 1 | 8, 17, 26 | 0.40, 0.40, 0.20 | 0.5495 |
|  | 2 | 9, 17, 22 | 0.45, 0.35, 0.20 | 0.5415 |
|  | 3 | 9, 16, 22 | 0.40, 0.30, 0.30 | 0.5308 |
| o__Cyanobacteriales (410/585) | 1 | 8, 10, 20 | 0.05, 0.20, 0.75 | 0.8240 |
|  | 2 | 9, 10, 20 | 0.05, 0.20, 0.75 | 0.8154 |
|  | 3 | 8, 10, 20 | 0.05, 0.25, 0.70 | 0.8199 |
| c__Dethiobacteria (72/102) | 1 | 20 | 1.00 | 0.6981 |
|  | 2 | 21, 22 | 0.40, 0.60 | 0.6890 |
|  | 3 | 20, 22, 34 | 0.40, 0.50, 0.10 | 0.7140 |
| f__Neisseriaceae (102/145) | 1 | 9, 18, 19 | 0.15, 0.65, 0.20 | 0.9052 |
|  | 2 | 10, 19 | 0.15, 0.85 | 0.9069 |
|  | 3 | 6, 10, 19 | 0.05, 0.20, 0.75 | 0.9082 |

### 2.5 Ok-*k*-mer runtime

Global search finds the highest-scoring ensemble within the user-defined length range and weight grid, but evaluating all candidates can be computationally expensive, particularly for larger datasets. To reduce search time, Ok-*k*-mer also provides a local search that starts from the best single *k*-mer length and expands the ensemble by varying previously selected lengths within a neighborhood of *±*Δ_*k*_ while adding a length from the full allowed range. We compared local search with global search on fullset Dashing2 distances across eight datasets (Methods 4.5 and 4.6). Ok-*k*-mer local search achieved a 1.15–6.41*×* speedup over global search (Table 3). Caldisericia showed the smallest speedup; both modes stopped after evaluating ensembles of up to two lengths because adding a second length did not improve correlation. Peak resident memory remained below 0.4 GiB for both modes across all eight datasets; local search used approximately 0.5–5.7% less memory than global search. Runtime and peak memory usage for the initial Dashing2 distance calculations are reported separately in Table A2.

**Table 3.**
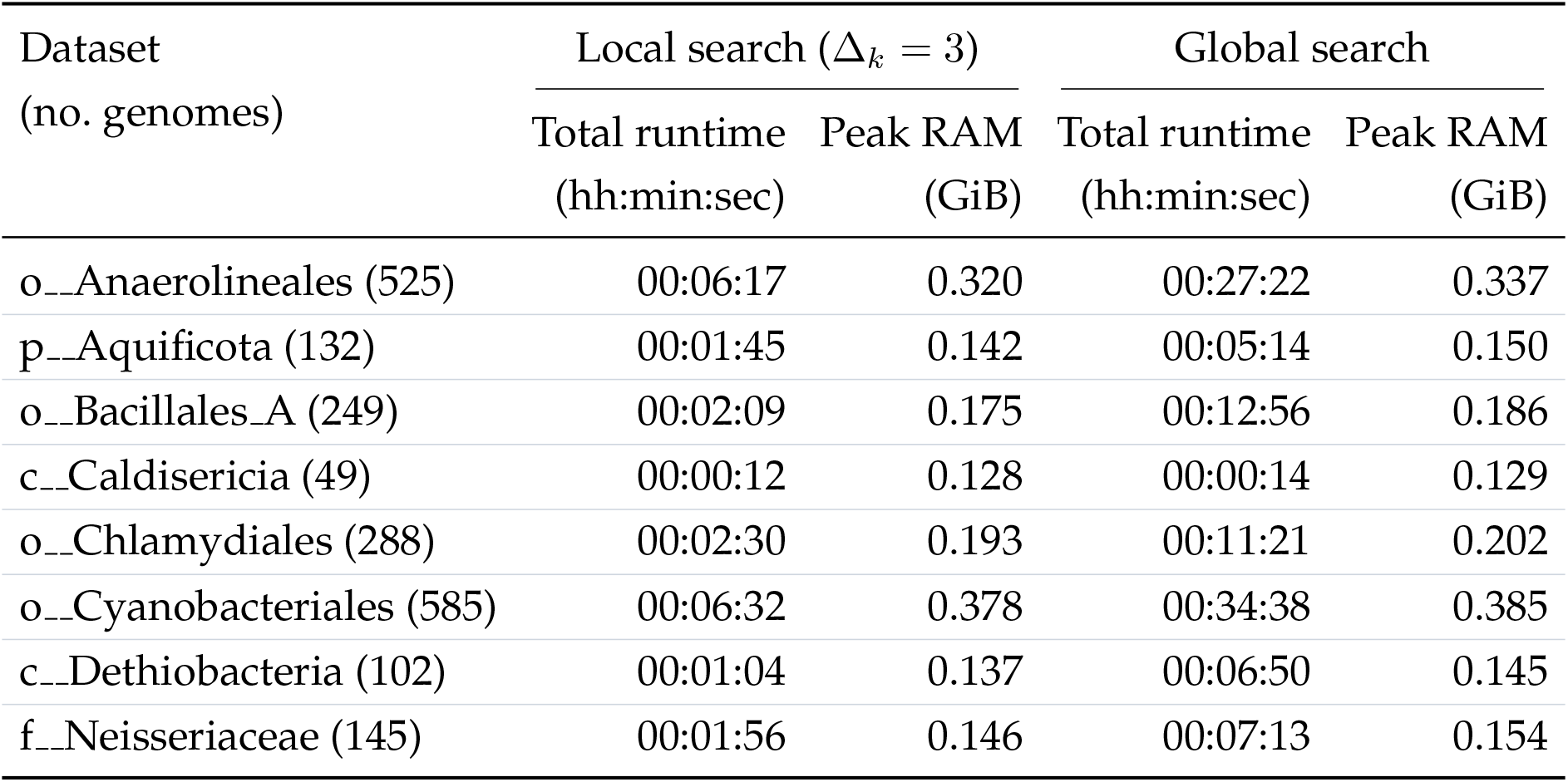
Runtime and peak memory usage for Ok-*k*-mer searches using full-set (unsketched) Dashing2 distances, measured with GNU `/usr/bin/time`. Each search used eight threads, considered up to three *k*-mer lengths from *k* ∈ {6, … , 40} with early stopping, and evaluated weights in increments of 0.05. Local search used Δ_*k*_ = 3. Total runtime is the elapsed wall time, including index loading and result writing; installation, index preparation, and Dashing2 computation are excluded. Peak RAM is the maximum resident set size in GiB.

| Dataset<br>(no. genomes) | Local search (Δ<sub>k</sub> = 3) |  | Global search |  |
|---|---|---|---|---|
|  | Total runtime<br>(hh:min:sec) | Peak RAM<br>(GiB) | Total runtime<br>(hh:min:sec) | Peak RAM<br>(GiB) |
| o__Anaerolineales (525) | 00:06:17 | 0.320 | 00:27:22 | 0.337 |
| p__Aquificota (132) | 00:01:45 | 0.142 | 00:05:14 | 0.150 |
| o__Bacillales_A (249) | 00:02:09 | 0.175 | 00:12:56 | 0.186 |
| c__Caldisericia (49) | 00:00:12 | 0.128 | 00:00:14 | 0.129 |
| o__Chlamydiales (288) | 00:02:30 | 0.193 | 00:11:21 | 0.202 |
| o__Cyanobacteriales (585) | 00:06:32 | 0.378 | 00:34:38 | 0.385 |
| c__Dethiobacteria (102) | 00:01:04 | 0.137 | 00:06:50 | 0.145 |
| f__Neisseriaceae (145) | 00:01:56 | 0.146 | 00:07:13 | 0.154 |

### 3 Conclusion

A major challenge for *k*-mer-based approaches going forward is to establish a rigorous way for users to select the value or values of *k* that should be used in a given situation. Here we introduced Ok-*k*-mer as a principled strategy for selecting either a single *k*-mer length or a multi-*k*-mer length ensemble that is well suited to computing whole-genome similarities across a phylogenetic tree. Across eight example GTDB datasets, we found that selecting and combining multiple *k*-mer lengths improves agreement with reference phylogenetic distances. Length selection depends on the dataset; different datasets favor different *k*-mer ensembles (Table 1). Of the datasets, Chlamydiales demonstrated the greatest improvement, an approximately 24% increase in Spearman *ρ* over the best single length using an ensemble of three *k*-mer lengths. Some datasets do well with a single dataset-specific *k*-mer length, but selecting this “best” *k*-mer length with Ok-*k*-mer still provides an improvement over a naive default *k*-mer length. Compared with ensembles recommended by Ok-*k*-mer, fixed *k* = 31 yielded correlations averaging 10.33% lower.

Subtree annotations provide a practical basis for choosing lengths for individual subtrees. Ok-*k*-mer performance remains stable under genome subsampling; across masking fractions up to 60%, mean correlations remain broadly consistent within each dataset. While users may not always have a full phylogenetic tree, the stability results demonstrate that we learn dataset-specific information that can generalize to larger sets of the same taxonomy. Given an incomplete reference tree, a potential application is to place an unknown genome using the selected *k*-mer recommendations. We provide multiple resolutions for our *k*-mer search: a single-best *k*-mer search and global and local searches for multiple *k*-mers. Additionally, while we use Dashing2 without sketching for the most accurate results, other *k*-mer-based initial pairwise distance approaches work. Using Dashing2 with sketching reduced the runtime of the initial pairwise distance calculations by approximately 28–754*×* across datasets relative to full-set calculations (Table A2), but yielded variable *k*-mer length ensembles. Future work will evaluate the tradeoff between sketch size, computational cost, and accuracy.

Ok-*k*-mer comes with some limitations that necessitate further investigation. The current objective requires a starting set of reference pairwise phylogenetic distances. Without these distances, we cannot determine the ranking of genome pairs. In the future it will be important to generalize Ok-*k*-mer to a flat set of genomes, without needing foreknowledge of an initial set of pairwise distances. Another major area for future work is to better understand why specific *k*-mer lengths are chosen. Based on the *k*-mer ensembles we have calculated to date, we contend that the distinct *k*-mer lengths chosen tend to capture complementary information. We have two hypotheses: (1) the *k*-mer choices arise from different mixtures of genomes in the dataset, and (2) the *k*-mer choices capture information from different compartments of a genome. The first hypothesis could explain why merging subtrees could result in varying *k*-mer choices. The second hypothesis is motivated by differences in selective pressures and sequence conservation between core and accessory regions, which may favor different *k*-mer lengths.

## 4 Methods

### 4.1 Evaluation datasets

The Genome Taxonomy Database (GTDB) provides reference phylogenetic trees with branch lengths [14]. Following the datasets analyzed in EvANI [2], we evaluated Ok-*k*-mer on eight clades from GTDB release R220 (R09-RS220 / 220.0). These datasets span bacterial taxonomic ranks from family to phylum and range from 49 to 585 genomes: o Anaerolineales, p Aquificota, o Bacillales A, c Caldisericia, o Chlamydiales, o Cyanobacteriales, c Dethiobacteria, and f Neisseriaceae. Dataset name prefixes indicate taxonomic rank. The genomes were downloaded individually from NCBI Assembly using GenBank FASTA URLs.

### 4.2 EvANI benchmarking framework for distance estimation

The EvANI benchmarking framework [2] takes both input genome sequences and corresponding reference phylogenies. It then evaluates various distance estimation tools by measuring the strength of the rank correlation between the tool’s distance estimates and the true pairwise tree distances. A higher Spearman correlation *ρ* between the ranks of computed genomic distances and reference tree distances indicates a more accurate distance measure, i.e. one that more accurately recapitulates true tree distances. We adopt this same framework since our objective is also to preserve the ordering of pairwise phylogenetic distances without assuming a linear relationship between genomic and phylogenetic distances.

### 4.3 Pairwise *k*-mer similarity calculation

For a fixed *k*-mer length *k*, let 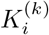 and 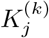 denote the sets of distinct *k*-mers in genomes *G*_*i*_ and *G*_*j*_, respectively. The Jaccard index is 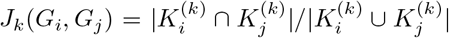, the fraction of distinct *k*-mers shared by the two genomes among those present in either genome [4]. It ranges from 0 for disjoint sets to 1 for identical sets and ignores *k*-mer multiplicities. The corresponding Jaccard distance is *d*_*J,k*_(*G*_*i*_, *G*_*j*_) = 1 − *J*_*k*_(*G*_*i*_, *G*_*j*_), so larger values indicate greater dissimilarity. While we use Dashing2, alternative *k*-mer counters, like KMC3, may be used to compute the distance between genomes [4, 15, 16].

### 4.4 Runtime and memory measurements

We use Dashing2 [17], with no sketching, to compute the initial all-pairs Jaccard distance with all *k*- mers, *k* ∈ {6, … , 40}. Dashing2 uses an underlying SetSketch data structure to count *k*-mers. dashing2 d i s t −− **s e t** −−long −kmers −k “ $k” −p *<*threads *>* −−square \ −F genome paths . t x t *>* “ j a c c a r d k $ {k} . t sv “

We ran Dashing2 release `v2.1.21` on all datasets with eight threads (`-p 8`) and recorded elapsed wall time and peak resident memory with GNU `/usr/bin/time`. For the sketch-based runs, we replaced `--set` with `--full -S 4096`, using full SetSketch with 4,096 registers and retaining `--long-kmers`.

We separately measured Ok-*k*-mer search runtime and peak resident memory for local (Δ_*k*_ = 3) and global searches using full-set and sketch-based distances (Tables 3 and A3). These measurements used eight search threads and considered up to three lengths with normal early stopping, terminating when an additional length failed to improve the best *ρ*.

### 4.5 Ok-*k*-mer search framework and global search

Ok-*k*-mer progressively selects *k*-mer lengths and weights to maximize concordance with a reference phylogeny. For an ensemble of *m k*-mer lengths, let *r*_*k*_(*p*) denote the rank of the Jaccard distance for genome pair *p* among all genome pairs at *k*-mer length *k*. We rank the reference tree distances once and separately rank the Jaccard distances for each *k*-mer length *k* ∈ {*k*_low_, … , *k*_high_} (Methods 4.3). By default, *k*_low_ = 6 and *k*_high_ = 40. We compute the weighted-average rank score 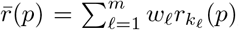, where weights are strictly positive and evaluated in increments of *w*_step_ (default: 0.05), subject to 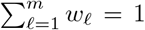 . Consistent with the EvANI benchmarking framework (Methods 4.2), we score each *k*- mer length ensemble by its Spearman correlation *ρ* calculated between 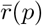 and reference tree distances across genome pairs.

The Ok-*k*-mer search proceeds from one length to the user-specified maximum ensemble size, optimizing weights for every candidate combination and retaining the *k*-mer lengths and weights achieving the highest *ρ*. The search stops when the best correlation at the next ensemble size fails to improve the previous best *ρ*, or when the maximum ensemble size is reached. Ok-*k*-mer then returns the best retained ensemble. At each evaluated ensemble size, Ok-*k*-mer global search evaluates every combination of distinct lengths within the allowed *k*-mer range and weight combination.

### 4.6 Ok-*k*-mer local search

Ok-*k*-mer local search uses the same scoring objective, weight grid, and stopping rule as global search, but restricts the candidate length combinations. The local search first evaluates every *k*-mer length in the user-specified range. This stage selects the best single *k*-mer length. At each subsequent stage, local search jointly evaluates combinations of one additional length from the full user-specified range and replacements within *±*Δ_*k*_ of each previously selected *k*-mer length. By default, Δ_*k*_ = 3, and candidate replacement lengths are restricted to {*k*_low_, … , *k*_high_}.

For example, suppose the allowed range is *k* ∈ {6, … , 40}, Δ_*k*_ = 3, and the best two-length ensemble is (20, 34). The three-length local search then evaluates combinations formed from one length in 17–23, the second in 31–37, and the third anywhere in 6–40. Both previously selected lengths can therefore change, while the additional length is unrestricted within the allowed range. This restriction reduces the candidate search space but can exclude the best ensemble found by Ok-*k*-mer global search at a given size.

## Supplementary information

The Ok-*k*-mer software is available at: https://github.com/StephenHwang/Ok-k-mer.

Code for the experiments presented in this manuscript is available at: https://github.com/StephenHwang/Ok-k-mer Experiments.

## Acknowledgements

We thank Sina Majidian for his work on the related EvANI project and his initial input to this project.

## Appendix A Supplementary Information

**Table A1.** Selected *k*-mer-based tools, their default *k*-mer lengths, and the role of *k*-mers in each method. Defaults serve different tasks, including whole-genome comparison, local mapping, and composition-vector analysis, and should be interpreted accordingly. The shorter default *k*-mer lengths used by FastANI and skani support local mapping and homologous-region detection for ANI estimation, rather than direct comparison of whole-genome *k*-mer sets.

| Tool | Purpose | Default *k* | Note |
|---|---|---|---|
| Mash [4] | Estimates whole-genome distance. | *k* = 21 | Uses MinHash sketches of whole-genome *k*-mers to estimate mutation distance without alignment. |
| sourmash [5] | Estimates similarity, containment, and ANI. | *k* = 31 | Uses scaled *k*-mer sketches for similarity and containment comparisons between unequal-sized inputs. |
| Skmer [6] | Estimates genomic distance from low-coverage, unassembled reads. | *k* = 31 | Compares read-derived *k*-mer sketches, correcting for incomplete coverage and sequencing error. |
| Dashing 2 [17] | Estimates Jaccard similarity, containment, and ANI. | *k* = 31 | Uses SetSketch for *k*-mer set overlap; weighted modes account for multiplicities. |
| Kmer-db [7] | Computes pairwise genomic distances using a reusable database. | *k* = 18 | Counts shared distinct *k*-mers across whole inputs, with optional MinHash subsampling. |
| FastANI [8] | Estimates ANI between genome assemblies. | *k* = 16 | Maps 3-kb fragments with MashMap's *k*-mer sketches; averages identity across reciprocal mappings. |
| skani [9] | Estimates ANI and aligned fraction. | *k* = 15 | Chains positional *k*-mer matches to identify homologous regions; estimates ANI from local *k*-mer containment. |
| AAF [18] | Constructs phylogenetic distances and trees from unassembled reads. | *k* = 25 | Uses read-derived *k*-mer presence/absence, with corrections for sequencing error and incomplete coverage. |

**Table A2.** Runtime and peak memory usage for full-set and sketch-based Dashing2 pairwise similarity calculations as measured with GNU `/usr/bin/time`. Each *k*-mer length (*k* = 6, … , 40) was run separately with eight Dashing2 threads. Total runtime is the sum of elapsed wall times across all *k*-mer lengths in hours:minutes:seconds. Peak RAM is the maximum peak resident memory across all runs. Mean absolute error (MAE) is the mean of the per-*k* mean absolute errors of sketch-based estimates relative to full-set results.

| Dataset | Dashing2 full |  | Dashing2 sketching |  | MAE |
|---|---|---|---|---|---|
|  | Total runtime<br>(hh:min:sec) | Peak RAM<br>(GiB) | Total runtime<br>(hh:min:sec) | Peak RAM<br>(GiB) |  |
| o__Anaerolineales (525) | 32:58:59 | 4.346 | 00:08:32 | 0.065 | 0.000791 |
| p__Aquificota (132) | 00:58:29 | 1.524 | 00:01:11 | 0.032 | 0.000957 |
| o__Bacillales_A (249) | 08:02:47 | 3.819 | 00:04:29 | 0.071 | 0.000965 |
| c__Caldisericia (49) | 00:12:36 | 1.512 | 00:00:27 | 0.019 | 0.000919 |
| o__Chlamydiales (288) | 05:01:16 | 2.348 | 00:02:54 | 0.042 | 0.000779 |
| o__Cyanobacteriales (585) | 212:28:13 | 7.395 | 00:16:55 | 0.119 | 0.000959 |
| c__Dethiobacteria (102) | 00:53:42 | 2.138 | 00:01:09 | 0.019 | 0.000843 |
| f__Neisseriaceae (145) | 01:46:18 | 2.019 | 00:01:48 | 0.041 | 0.001282 |

**Table A3.** Runtime and peak memory usage for Ok-*k*-mer searches using sketch-based Dashing2 distances, measured with GNU `/usr/bin/time`. Dashing2 used full SetSketch with 4,096 registers (`--full -S 4096`). Each search used eight threads, considered up to three *k*-mer lengths from *k* ∈ {6, … , 40} with normal early stopping, and evaluated weights in increments of 0.05. Local search used Δ_*k*_ = 3. Total runtime is the elapsed wall time, including index loading and result writing; installation, index preparation, and Dashing2 computation are excluded. Peak RAM is the maximum resident set size in GiB.

| Dataset<br>(no. genomes) | Local search (Δ<sub>*k*</sub> = 3) |  | Global search |  |
|---|---|---|---|---|
|  | Total runtime<br>(hh:min:sec) | Peak RAM<br>(GiB) | Total runtime<br>(hh:min:sec) | Peak RAM<br>(GiB) |
| o__Anaerolineales (525) | 00:04:17 | 0.305 | 00:22:29 | 0.312 |
| p__Aquificota (132) | 00:01:07 | 0.140 | 00:08:56 | 0.151 |
| o__Bacillales_A (249) | 00:02:37 | 0.171 | 00:10:24 | 0.184 |
| c__Caldisericia (49) | 00:01:08 | 0.128 | 00:05:15 | 0.137 |
| o__Chlamydiales (288) | 00:02:26 | 0.188 | 00:09:49 | 0.198 |
| o__Cyanobacteriales (585) | 00:07:09 | 0.347 | 00:30:52 | 0.360 |
| c__Dethiobacteria (102) | 00:01:02 | 0.135 | 00:06:35 | 0.147 |
| f__Neisseriaceae (145) | 00:01:37 | 0.146 | 00:07:19 | 0.154 |

**Table A4.** Comparison of fixed single-value 21-mer and 31-mer correlations with the optimal Ok-*k*-mer recommendation across GTDB datasets. Percentages in parentheses show the relative change in *ρ* compared with the corresponding Ok-*k*-mer result.

| GTDB Dataset<br>(no. genomes) | *k* = 21 | *k* = 31 | Ok-*k*-mer |  |
|---|---|---|---|---|
|  | ρ<br>(% change) | ρ<br>(% change) | *k*<sub>1</sub>, …, *k*<sub>*n*</sub><br>(*w*<sub>1</sub>, …, *w*<sub>*n*</sub>) | ρ |
| o__Anaerolineales (525) | 0.6230<br>(−1.51%) | 0.5596<br>(−11.54%) | 10, 20, 39<br>(0.15, 0.80, 0.05) | 0.6326 |
| p__Aquificota (132) | 0.6873<br>(−0.85%) | 0.6715<br>(−3.13%) | 19, 20, 34<br>(0.35, 0.40, 0.25) | 0.6932 |
| o__Bacillales_A (249) | 0.6922<br>(−1.03%) | 0.6756<br>(−3.40%) | 7, 23, 24<br>(0.25, 0.40, 0.35) | 0.6994 |
| c__Caldisericia (49) | 0.7660<br>(−0.94%) | 0.6893<br>(−10.86%) | 20<br>(1.00) | 0.7733 |
| o__Chlamydiales (288) | 0.4188<br>(−20.29%) | 0.3843<br>(−26.86%) | 9, 17, 22<br>(0.45, 0.30, 0.25) | 0.5254 |
| o__Cyanobacteriales (585) | 0.7900<br>(−3.22%) | 0.6896<br>(−15.52%) | 9, 10, 20<br>(0.05, 0.20, 0.75) | 0.8163 |
| c__Dethiobacteria (102) | 0.7005<br>(−0.06%) | 0.6577<br>(−6.16%) | 20, 22<br>(0.70, 0.30) | 0.7009 |
| f__Neisseriaceae (145) | 0.8952<br>(−1.66%) | 0.8630<br>(−5.20%) | 10, 19<br>(0.20, 0.80) | 0.9103 |

**Fig. A1.**
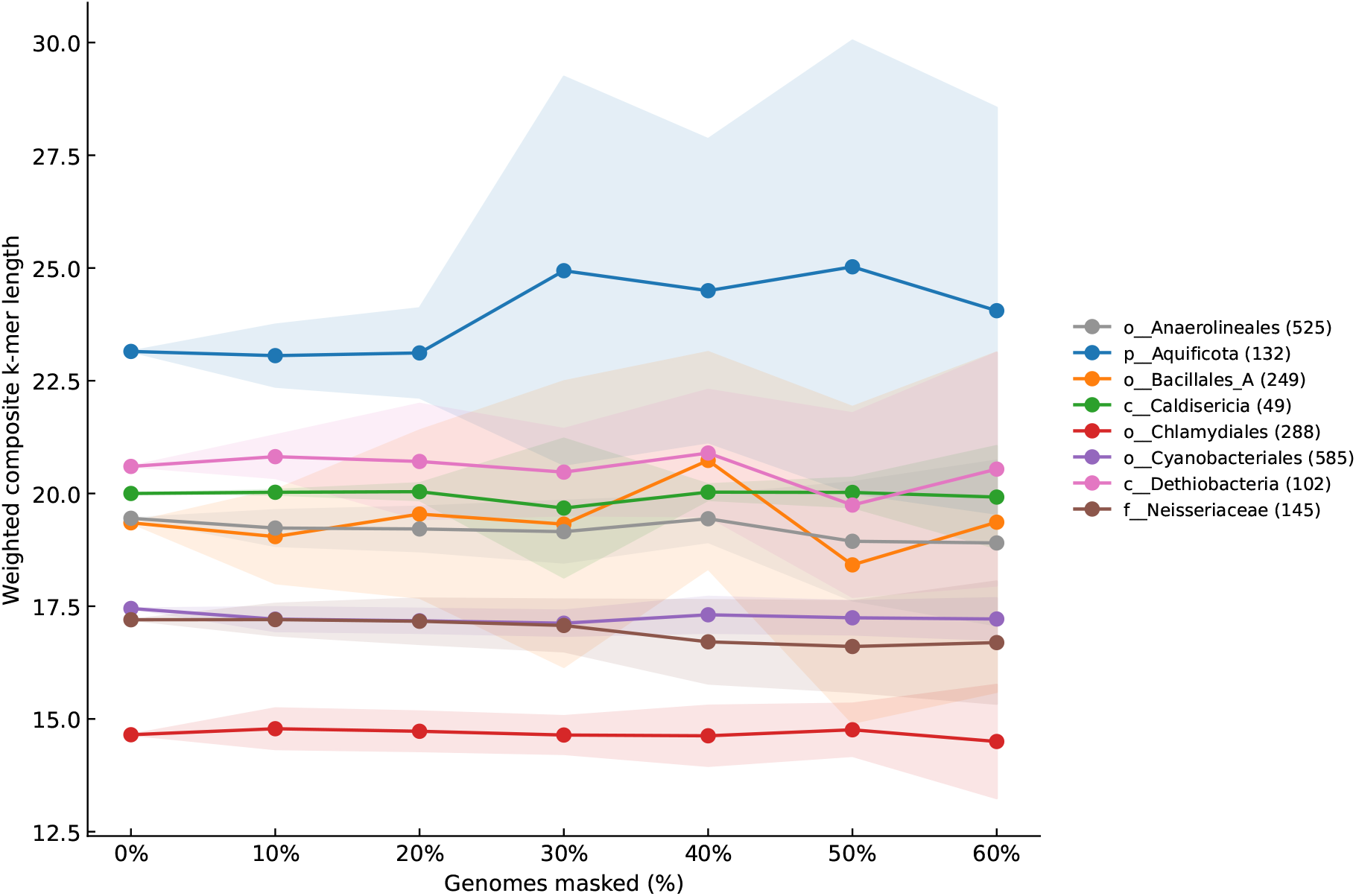
Weighted composite ranks of Ok-*k*-mer *k*-mer recommendations across eight GTDB datasets at masking fractions from 0% to 60% in 10% increments. The weighted composite rank is the sum of the *k*-mer lengths multiplied by their respective weights. The spread is *±*1 standard deviation of the weighted composite rank.

